# A fuzzy-registration approach to track cell divisions in time-lapse fluorescence microscopy

**DOI:** 10.1101/419572

**Authors:** Saoirse Amarteifio, Todd Fallesen, Gunnar Pruessner, Giovanni Sena

## Abstract

**Background:** Particle-tracking in 3D is an indispensable computational tool to extract critical information on dynamical processes from raw time-lapse imaging. This is particularly true with *in vivo* time-lapse fluorescence imaging in cell and developmental biology, where complex dynamics are observed at high temporal resolution. Common tracking algorithms used with time-lapse data in fluorescence microscopy typically assume a continuous signal where background, recognisable keypoints and independently moving objects of interest are permanently visible. Under these conditions, simple registration and identity management algorithms can track the objects of interest over time. In contrast, here we consider the case of transient signals and objects whose movements are constrained within a tissue, where standard algorithms fail to provide robust tracking.

**Results:** To optimize 3D tracking in these conditions, we propose the merging of registration and tracking tasks into a fuzzy registration algorithm to solve the identity management problem. We describe the design and application of such an algorithm, illustrated in the domain of plant biology, and make it available as an open-source software implementation. The algorithm is tested on mitotic events in 4D data-sets obtained with light-sheet fluorescence microscopy on growing *Arabidopsis thaliana* roots expressing CYCB::GFP. We validate the method by comparing the algorithm performance against both surrogate data and manual tracking.

**Conclusion:** This method fills a gap in existing tracking techniques, following mitotic events in challenging data-sets using transient fluorescent markers in unregistered images.

## 1 Background

It is generally understood that automated imaging and tracking methods can exceed manual tracking of objects in large data-sets. These solutions are of growing importance in life science applications, where time-lapse microscopy is a powerful and popular tool for capturing dynamics at cellular and sub-cellular scales [1]. Indeed, a wide range of microscopy [2] and computational approaches [3, 4, 5, 6] have been developed and adapted to particle tracking in biology. Developmental biology has benefit enormously from automated *in vivo* tracking methods, ranging from cell lineage tracing in both animals [7, 8, 9] and plants [10, 11, 12], to cell shape tracking [13, 14] and tracing of growing organs as in the case of plant roots [11, 15, 16, 17, 18].

It should be noted that in all these cases the signal to track is approximately constant, which generally makes the task easier notwithstanding occlusions or other objects leaving and entering the field of view. Instead, it is significantly more challenging to track an intermittent or transient signal, as in the case of cell divisions or other short-lived events [19, 20, 21], or a signal distributed in 3D space like in confocal or light-sheet microscopy stacks.

From a computational point of view, tracking in time-lapse image data involves detection of objects in individual frames and solving their identification across frames (identity management). The goal of identity management is to link an object in one frame to what is found to be the same object in a later frame, so that linked objects share the same identity label, or identifier. The set of linked identifications of a given object through time is referred to as its lineage.

Tracking algorithms are greatly influenced by the structure of the data. Macroscopic objects may have complex morphologies that can be used to improve their identification [22]. Objects smaller than imaging resolution, on the other hand, will appear as featureless blobs; in these cases tracking relies more strongly on dynamical models alone [23]. For example, “multiple-hypothesis tracking” [24, 25] takes an exhaustive and deterministic approach to consider all possible lineage trees. A similar approach has been previously applied to tracking plant root cell nuclei in 3D [11]. Stochastic algorithms have also been proposed [26], specifically to deal with particularly noisy data-sets. Whenever persistent features, or reference points known as keypoints, can be easily identified in the background, a common strategy is to build a link of the same keypoint in two frames [27, 28, 29, 30]. Keypoints appearing in more than one frame are defined as inliers, while those appearing in only one frame are treated as outliers. If inliers exist, a geometric transformation (including for example translation, rotation, reflection, scaling) linking them in the background can be determined and used as a first guess for tracking the objects of interest in the foreground.

Unfortunately, tracking transient events in 3D is still challenging and imprecise especially in plant biology where soft tissue rarely offers usable reference points. Here, we offer a novel solution to this problem, by exchanging the common method of morphological analysis [20] for a fuzzy registration-tracking approach, where *fuzzy* indicates a random sampling of points. As a case-study, we collected and analysed a 4D data-set of cell division events in the *Arabidopsis* root meristem, throughout several consecutive days. This was achieved with time-lapse 3D scanning of growing transgenic roots expressing the fluorescent reporter CYCB::GFP, through a previously described light-sheet microscope setup [16]. The mitotic events are here identified by the short-lived CYCB::GFP signal, sparsely distributed in a soft tissue lacking fluorescent reference keypoints and generally difficult to identify due to low signal-to-noise ratio. This represents a very common set of circumstances in time-lapse imaging of plant tissues. Due to its general strategy and features, we believe that the methodology proposed might be readily applied to similar 4D data-sets collected from other tissues.

## 2 Results

### 2.1 Tracking Algorithm

#### 2.1.1 General strategy

The identification of inliers is the key challenge in the kind of experimental dataset that we are discussing. When single-particle tracking methods require preregistration, they may not be robust against lack of guaranteed inliers. On the other hand, when single-particle tracking does not need pre-registration, it is typically based on models of the objects’ motion, which are not optimal with transient objects and with low signal-to-noise ratio across extended image sequences.

We treated the position, appearance and disappearance of transient objects as a random spatial process. The frame-to-frame displacement of objects embedded in a tissue can be described as the effect of two random variables: (1) large-scale movements of the tissue within the field of view, due for example to its growth or to the inability of the microscope to focus on a fixed point of the field of view over time; (2) small-scale fluctuations of the objects of interests within the tissue.

The main purpose of the algorithm is to find the best large-scale transformation that explains most of the object correspondence, despite the noisy small-scale dynamics. The strategy adopted was to find correspondences between the random process at time *t* and the random process at time *t* – *τ* where *τ* is a lag variable. In this work, we only considered a *τ* = 1step, which is equivalent to approximating the tissue as effectively rigid at the given temporal scale. As mentioned above, objects that appear in both frames are called inliers and those that do not appear in both frames are called outliers and cannot be linked by a global frame-frame transformation. For example, outliers can be either debris, *i.e.* any object which is determined not to be an object of interest, or could be an object of interest that has just exited or entered the new frame.

There are two main steps in the execution of the algorithm: (i) candidate transformations are generated from the data; (ii) among these candidates, the optimal transformation is chosen with respect to some previously defined objective. Inliers are then mapped onto each other by the chosen global transformation and labelled (with identifiers). Finally, new identifiers are generated for the outliers within the region of interest.

Algorithm 1 provides a terse, high-level overview of the method. The key algorithm stages, which amount to finding the best global transformation from the data, are discussed below.

#### 2.1.2 Proposing transformations

Consider the sets of objects *U*(*t*),*V*(*t* – *τ*) at times offset by a lag *τ*. A subset of these corresponds to true objects of interest (*i.e.* not noise or debris) and a further subset of these exists in both frames. Given *N* objects at time *t* and *M* objects at time *t* — *τ*, a number of permutations 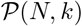 are sampled from *U* and a number of permutations 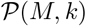 are sampled from *V*. We limited our method to *k* ∈ {1,3}, thus sampling either single objects (*k* = 1) or triangular *constellations* (*k* = 3) from each frame. Objects take a natural order and the same object can only appear in one constellation.

Let *n* be the number of *k*-constellations *c_i_*(*t*) sampled from *U*(*t*) and *m* be the number of *k*-constellations *c_j_*(*t*′ = *t* – *τ*) sampled from *V*(*t* – *τ*). In general, for two tensors **M**_1_ and **M**_2_ related by **M**_1_ = **AM**_2_, the affine transformation **A** can be uniquely determined through a least squares method (this is, in general, over-determined). Here, the transformation *γ_ij_* is the affine transformation, taking the set *c_i_*(*t*) to the set *c_j_* (*t*′): *c_j_* (*t*′) = ∑ *γ_ij_ c_i_*(*t*)

##### Algorithm 1 Transient point cloud tracking algorithm

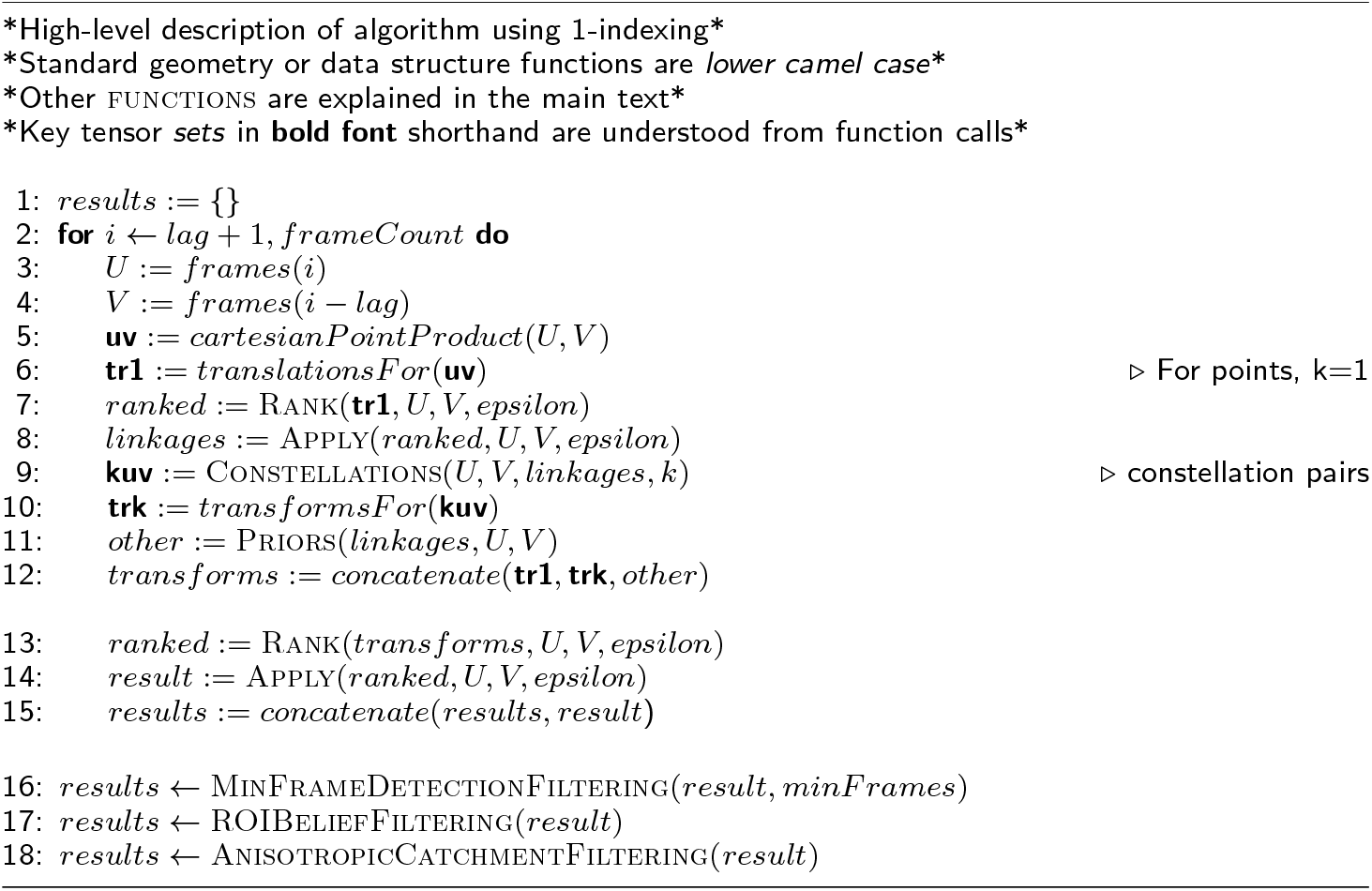

In lines 3-6 of the Algorithm 1, the constellation are sampled in frames *U* and *V*. Translations are then generated for each pair of objects *v* ∈ *V* and *u* ∈ U, and added to the list of candidate transformations.

Constellation sampling (line 9) is preferably seeded with linkages found by translations so that likely outliers are excluded from consideration. If a given translation maps *n* objects in *U* to *n* objects in *V*, then these object-object pairings can be considered candidate linkages. This not only reduces the candidate objects to those likely inliers, but constellation congruences can be identified as well, so the same set of objects in one frame can be paired to the same set of objects in the other frame when proposing transformations. Constellation pairs 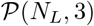, 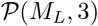 can then be sampled from the *L* inliers appearing in both frames. These will be used to generate proposal affine transformations, to provide a more optimal fit than translations alone.

#### 2.1.3 Ranking transformations

Transformation ranking was carried out using a modified least-squares-loss function. The least-squares-loss method is generally described as the minimization of 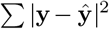, where **y** is a proposal vector and 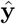 is the target vector. In our case, the proposal vectors are objects in the frame *U*(*t*) transformed back to *t*–*τ* by one of the proposed transformations. The target vectors are actual objects sampled in *V*(*t* – *τ*). The least-squares-loss objective was modified based on the following prior: (i) we expect a global rigid transformation to explain the movement of object centroids up to small fluctuations due to individual object movement which are deemed negligible. (ii) A distance > *ε* between objects’ centroids is assumed. The value of *ε* was determined from the data and is always slightly larger than the average object radius. When considering distances between proposal objects and target objects, only distances to the first nearest neighbour within a radial distance ε were considered. Each candidate transformation *γ* was applied to all objects *u_i_*(*t*) ∈ *U*(*t*), *i* ∈ {1, 2,…, *N*}. Let 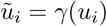. We used a cost function

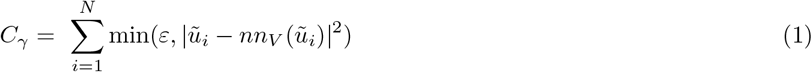

where 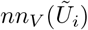 is the nearest neighbour position in *V*(*t*–*τ*) to the projected object 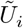. If the projected object had no nearest neighbour within a sphere of radius *ε*, the capped distance *ε* + 1 was attributed. This is illustrated in Fig. 1.

**Fig 1:**
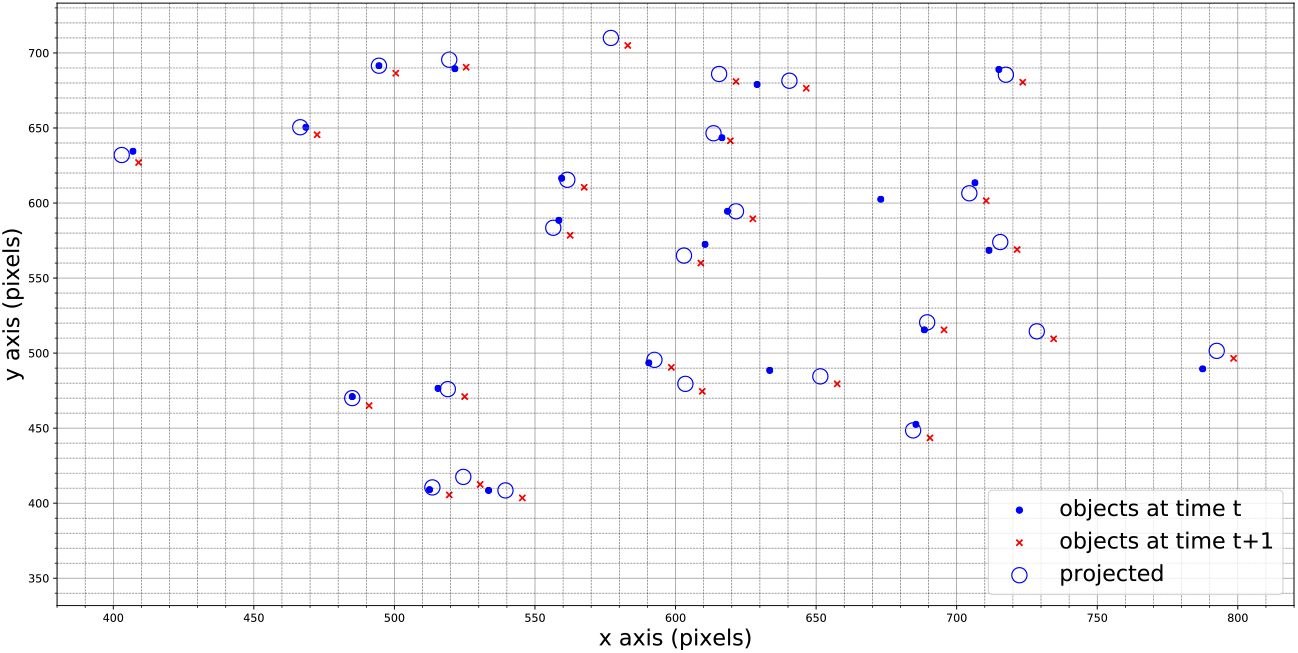
Evaluating transformations: red crosses correspond to objects in *V*(*t* – *τ*) and blue dots correspond to objects in *U*(*t*). The blue circles with radius *ε* correspond to the transformation of the set *U*(*t*) under the proposal transformation.

The cost function will rank transformations by how well they explain the movement of the majority of objects. If there are small differences in scores, we may prefer transformations that have the smallest displacement. For this reason, we can regularise this cost function to add a penalty to larger displacements.

### 2.2 Output: Life Matrix

The main output of the algorithm is a binary matrix called the Life Matrix, where each column corresponds to a time-point (i.e. a single frame in the time-lapse), each row contains the lifetime of a single detected object, and each element is equal to 1 if the object is detected in that frame, and to 0 otherwise. Debris exhibit movements that are not correlated with that of the objects of interest and therefore would typically appear in the Life Matrix as events with a life-span of a single frame. We discarded such single-frame events.

The algorithm was applied on time-lapse images of cell division events in roots of transgenic *Arabidopsis* plants expressing the fluorescent reporter CYCB::GFP, collected over a few days through light-sheet microscopy, following established methods [16]. A representative Life Matrix is shown in Fig. 2.

**Fig 2:**
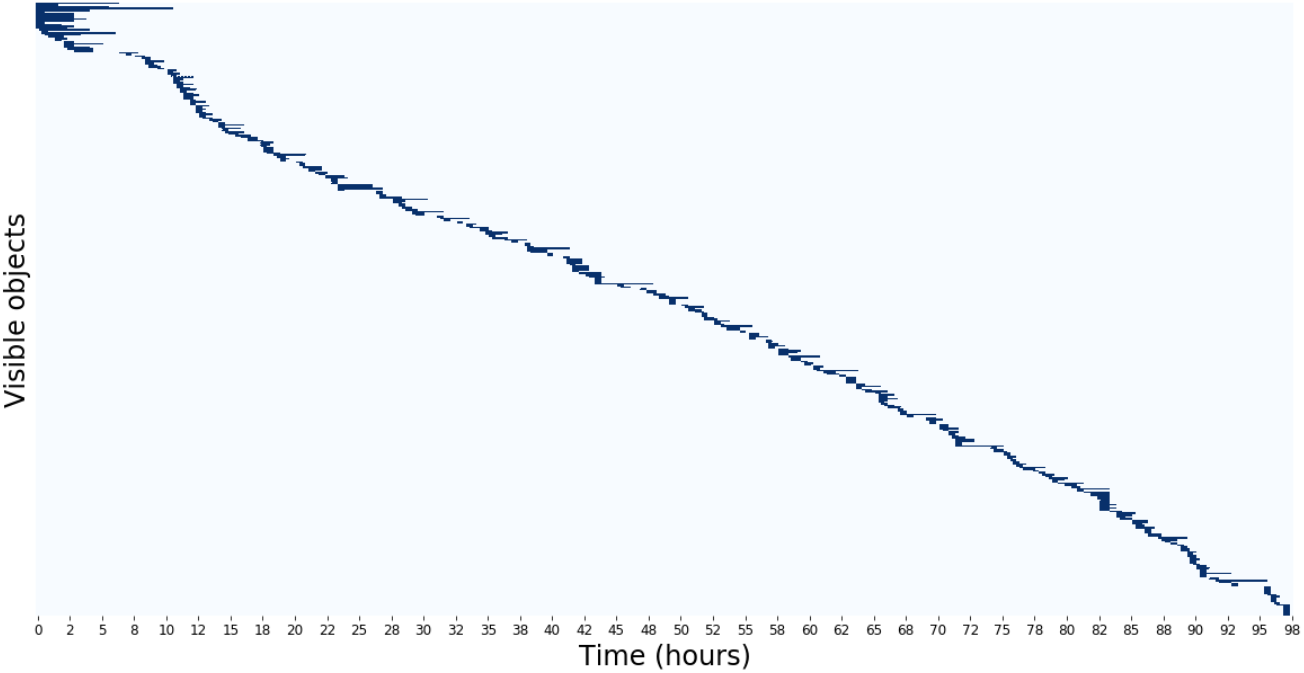
Life matrix of one representative root: each row represents the history of a single individual, detected (dark blue) at the corresponding time on the x-axis. The life matrix shows a generally consistent trend in the appearance and persistence of objects. At the beginning of this time-lapse, objects persisted for an unusual long time.

The number of objects observed as a function of time (Fig. 3, top) and the average duration of events (”mean age”, in number of frames) as a function of time (Fig. 3, bottom) are alternative representations of the data shown in the Life Matrix.

**Fig 3:**
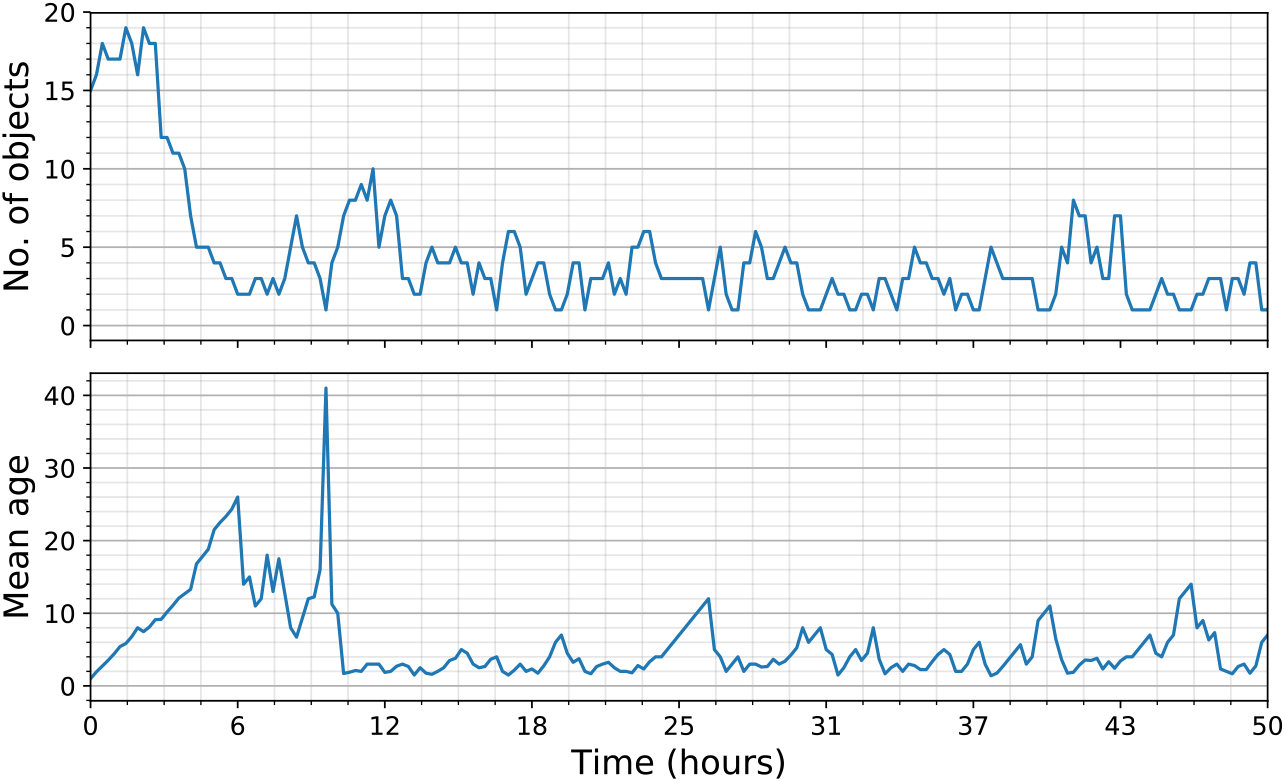
The population time series reflect the information shown in the life matrix described above. Top panel, number of events as a function of time; bottom panel, average event’s duration (in numbers of frames) as a function of time. The high spike in the mean age is a result of one object surviving for a long time while others disappear. After this point we observe more typical fluctuations in object activity.

### 2.3 Validation against surrogate data

We used computationally generated (surrogate) data to validate the tracking algorithm against a known spatial transformation. To be representative of the biological images intended to be tracked, the surrogate data were generated respecting a minimum object separation *ε*. A number of parameters were introduced (Table 1) to determine how data are generated and the tracker was tested for different points in the parameter space. The parameter *n* determined how many objects were randomly and uniformly distributed and added to the set of points *U*. A random Euclidean transformation (roto-translation) 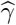 was generated using the parameters *r* and *s* to control the magnitudes of rotation and translation, respectively. A set *V* was produced by applying 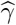 to *U*. The parameter *u* determined the strength of (small) random fluctuations to individual object locations in *V*, and *p* determined how many “noise” points were added to both *U* and *V* to represent debris or particles that existed in one but not the other frame. These added points also respected the minimum object separation *ε*. In the biological context this corresponds to cells that are no longer fluorescing or just beginning to fluoresce at time *t*.

**Table 1:** Surrogate data parameters: a random Euclidean transformation determined by parameters (*r, s, u*) is applied to a set of *n* objects *U* to produce a new set of objects *V*. To simulate noise, *p* objects are added at random locations to both *U* and *V*.

| Param. | Description |
| --- | --- |
| $\varepsilon$ | Minimum object separation |
| $n$ | Number of points to generate in the point-set $U$ |
| $p$ | Number of "noise" points to add to the point-set $U$ and the point-set $V$ |
| $r$ | Generate random (global) rotations in the range $[-\pi/r, \pi/r]$ |
| $s$ | Generate random (global) translations in the range $[0, s]$ |
| $u$ | Random uniform shift of points in $V$ on each axis in the range $[0, u]$ |

The tracking algorithm’s efficiency in finding the Euclidean transformation 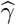 from *U* to *V* generally depends on the amount of noise and on the magnitude of the rotation and translation. To test our tracker’s performance, we established a classification task for the selected transformation: when an object in *U* was mapped onto a true object in *V*, it was considered an inlier and a value 0 was assigned, while a value −1 was assigned to outliers. A transformation’s score (or loss) was calculated as the sum of all these values and averaged over samples. The best transformation is the one with the lowest loss. This analysis showed that for small rotations (less than 5 degrees) or for small translations the tracker is robust and slowly degrades for larger rotations (more than 10 degrees), as shown in Fig 4.

**Fig 4:**
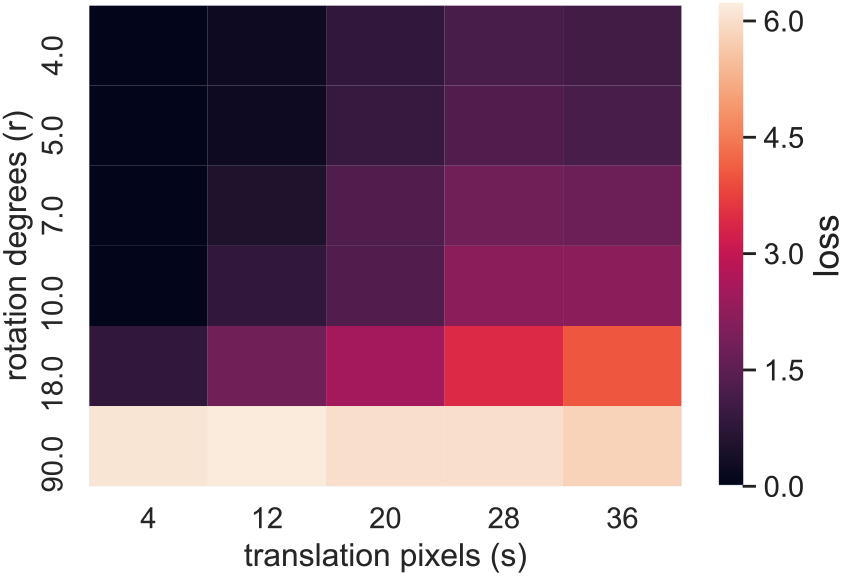
*Surrogate data results:* heat-map plot representing the scores of transformations with different parameters *s* and *r*.

### 2.4 Validation against manual tracking

The performance of the algorithm was also verified by manually marking the position of all GFP-expressing cells in a subset of 50 time-points by an experienced researcher without prior knowledge of the results of the algorithm output. Using the built-in FIJI plugin *CellCounter* the researcher manually identified 400 CYCB::GFP events which were present in at least 2 consecutive time-points, that is, using the same bias against single time-point events as the algorithm. In the same subset, the algorithm automatically detected 385 events. Given the 15-minute temporal resolution of our data-set and the longer persistence of the CYCB::GFP reporter, we reasoned that these should be considered either spurious events or plain artefacts. The rate of false negatives, *i.e.* events detected by the researcher but not by the algorithm, was 16.5 %, while the rate of false positives *i.e.* events detected by the algorithm but not by the researcher was 12.8 %. As a comparison, an analogous setup based on light-sheet microscopy had been previously used with *Arabidopsis* roots constitutively expressing the nuclear-localised, fluorescent fusion protein H2B::YFP [11]. In that study, the time-lapse imagine was performed with a temporal resolution comparable to the data presented here, but cell division events were identified as branching points on nuclei trajectories through a completely independent and competing approach. When validated against manual tracking, that method resulted in 26 % false negatives and 13 % false positives [11]. Given the fact that the goal is to extract statistical information from a limited sample of true events, it is generally preferable to minimize the false positives at the expense of the false negatives. In other words, it is acceptable to miss some true events because we are generally dealing with a sample anyway, but it should be avoided as much as possible to add spurious events to the sample. Taking this in consideration, we conclude that the method presented here is preferable when detecting cell divisions at high temporal resolution.

### 2.5 Validation with lag test

A third independent validation of our method was performed with a “lag test”. We counted the number of objects where identifiers were in agreement when performing the tracking using different lags *τ*. For example, we may expect the following equivalence

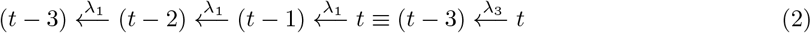

where, with an abuse of notation, *λ_τ_* corresponds to the highest-scoring transformation with lag *τ*. As a representative example of this test, we considered 734 objects in a sample sequence: 42 (or ≈ 6%) showed disagreements between λ_1_ and λ_2_ and, excluding these, 29 (or ≈ 4%) showed disagreements between λ_1_ and λ_3_ in specific frames (Figure 5).

**Fig 5:**
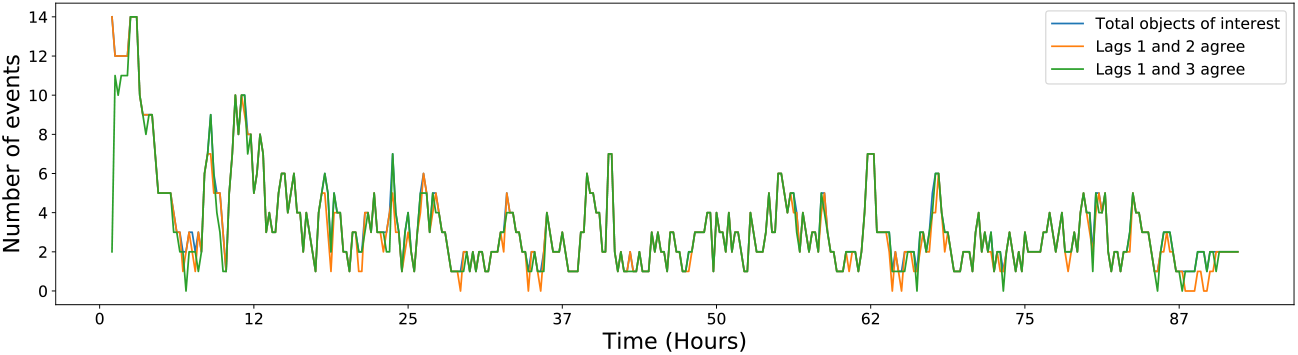
Lag test: identifiers for objects of interest (excluding single-frame detections) assigned by λ_1_ were compared with identifiers assigned by λ_2_ (orange line) and λ_3_ (green line) transformations.

### 2.6 Filtering

After tracking, filtering can be applied in a post-processing stage. A region of interest (ROI) was introduced to limit the analysis to the root meristem (ellipse in Fig. 6 and Fig. 7; see Methods). The colour code in Fig. 6 illustrates when and where events occur and indicates (roughly) if plotted points correspond to the same object. In Fig. 7, large angle differences between object displacement vectors and a global transformation vector appear in lighter colours (*e.g.* red) and are indicative of possible inlier mis-classification. Objects outside of (or far from) the ellipse and objects that show large angle differences are candidates for removal. In the case of angle differences, actual displacements and *ε* values can be taken into account when filtering objects. For example, in the final result output from the tracker, we could filter objects out of our final analysis when they met one of the following conditions: (i) the local displacement of an object away from the location predicted by the global transformation is above a certain threshold; (ii) the object is moving in a direction that is sufficiently different to the rest of the object population.

**Fig 6:**
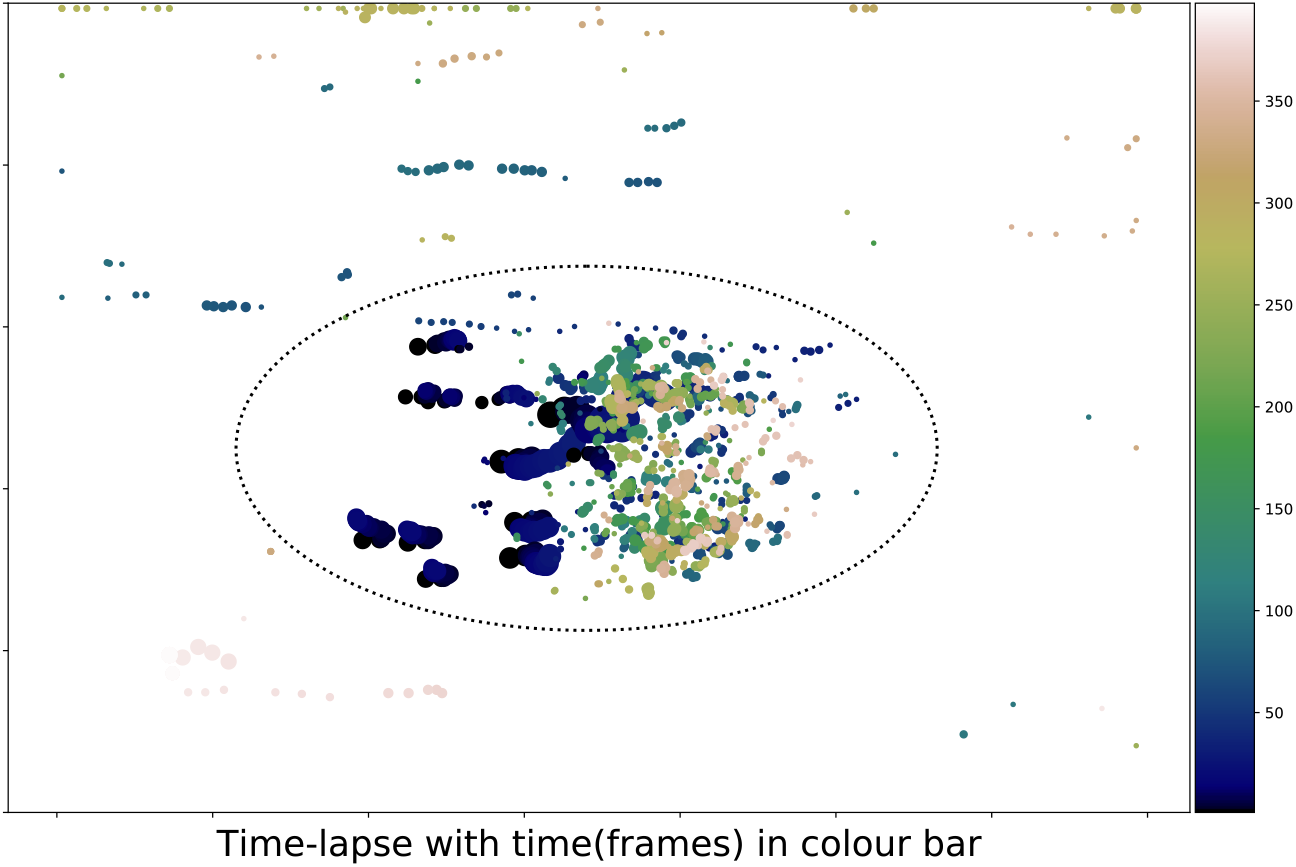
Time-lapse plots colour-coded for time. The ellipse represents the ROI over many frames.

**Fig 7:**
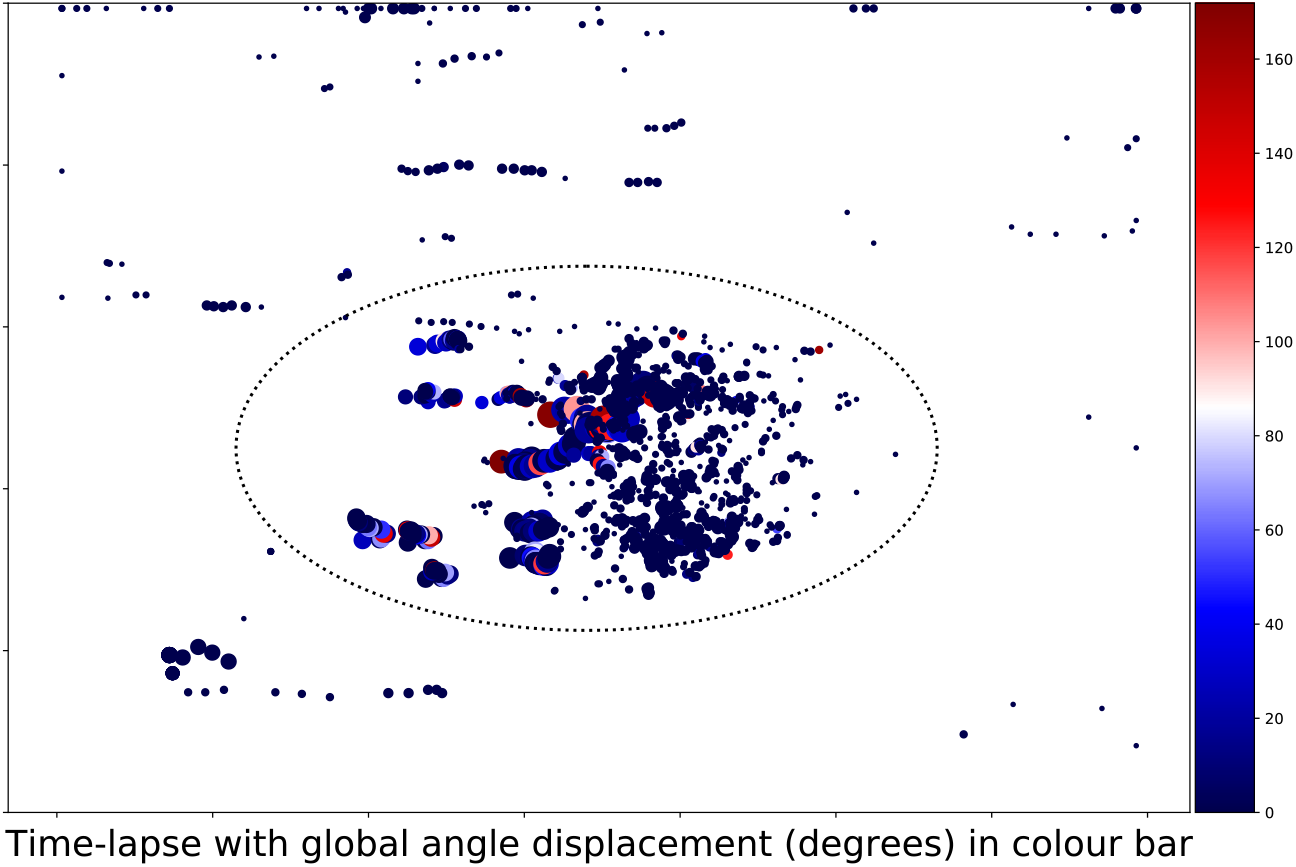
Time-lapse plots colour-coded for angle differences between the object displacement vector and the global transformation vector. The ellipse represents the ROI over many frames.

Fluctuations of the ROI area across frames are an indicator for how well the image processing step can identify the region of mitotic activity in the sample. When the signal-to-noise ratio is good, this region can be accurately detected and its relatively small area varies smoothly between frames. On the other hand, if there is too much noise or few true events, the identified ROI could become relatively large and vary non-smoothly from frame to frame. In the latter case, debris may lie within an excessively large ROI and be picked up by the tracker. For example, in Fig. 8a debris appear in the top part of the images, particularly towards the later stages of the experiment due to drift in microscope calibration or other factors. The root grows in the positive *x* direction (left to right) following the gravity vector, so precipitating debris appear moving in the same direction of root growth, while static marks or defects in the imaging chamber appear as moving in the opposite direction.

**Fig 8:**
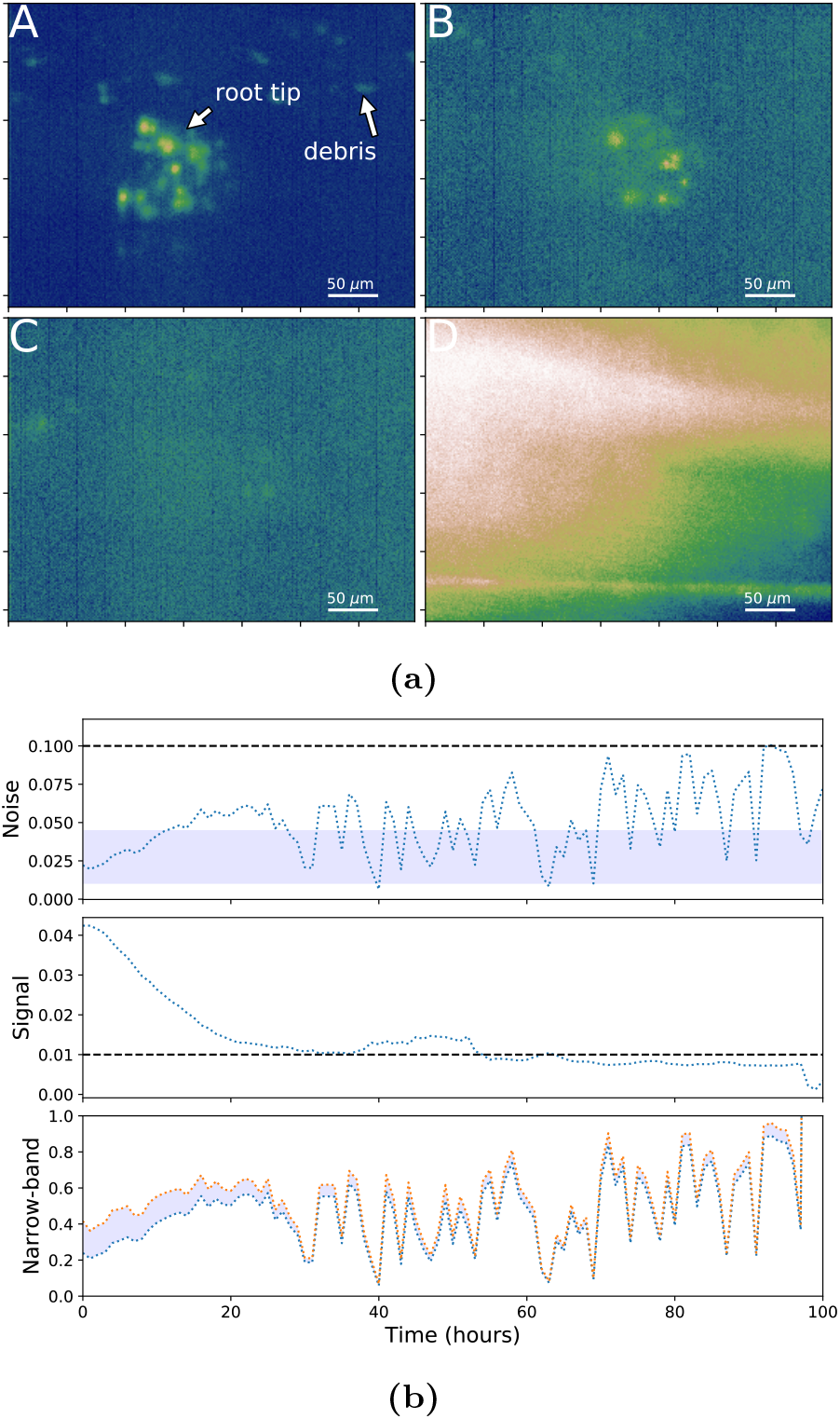
Variation in frame properties: the panels in (a) show four typical frames: A, “good” image; B, noisy image; C, noisy image with a low number of objects and some noise; D, over-saturated image to be marked as degenerate. The panels in (b) show a quantitative analysis over 100 consecutive sample frames from one representative root. The Signal and Noise levels are plotted with threshold values. The Narrow-band plot shows the region between the 95^*th*^ and 99^*th*^ percentile ranges of the image histogram. Notably, these values fluctuate dramatically.

### 2.7 Algorithm’s robustness and general performance

The described tracking algorithm generated comparable results when applied to different roots grown under the same conditions, showing expected robustness with respect to biological root-to-root variation. We sampled 200 frames each from 3 independent roots, and calculated a score for the displacement between predicted object locations and actual object locations: the three distributions showed no significant difference among them (Fig. 9). To measure the algorithm performance in dealing with variable object density, we plotted average and standard deviation of detection rates calculated from 200 frames, for three arbitrary values of *ε* (minimum required distance between distinct objects; see Methods).

**Fig 9:**
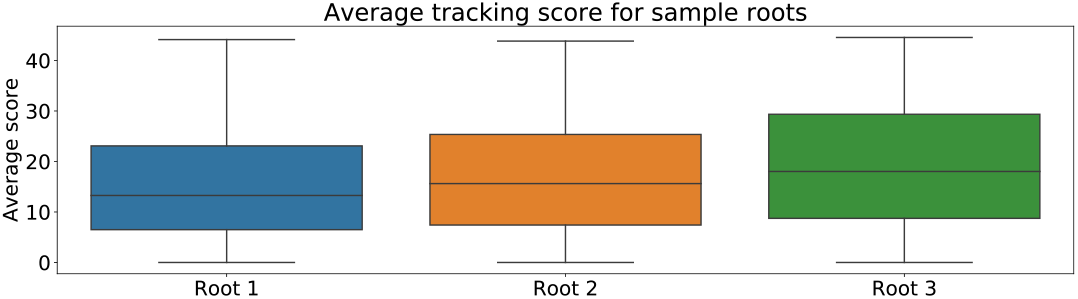
Performance for multiple root samples: box plots show the mean tracker score, averaged over 200 frames for each root.

Our results show that minimum performance is expected when the average number of objects per frame is between 5 and 10 (Fig. 10), while it increases rapidly for fewer and more objects per frame. Moreover, the variance in detection rate decreases dramatically when more than 15 objects per frame are present, as expected. with larger epsilon values (Fig. 11).

**Fig 10:**
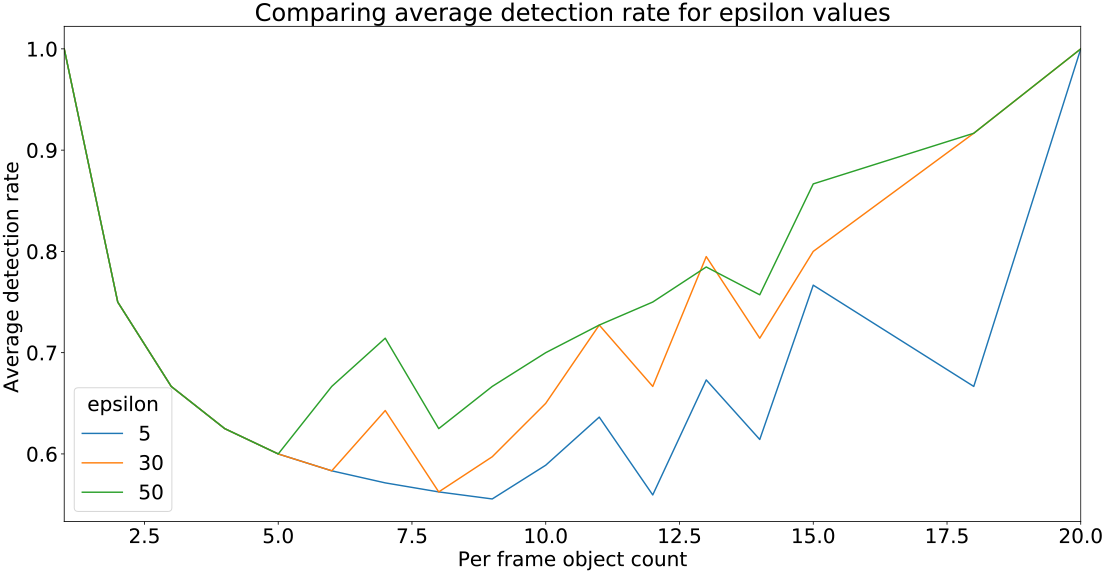
Average detection rate for 200 frames in each of the three sample roots.

**Fig 11:**
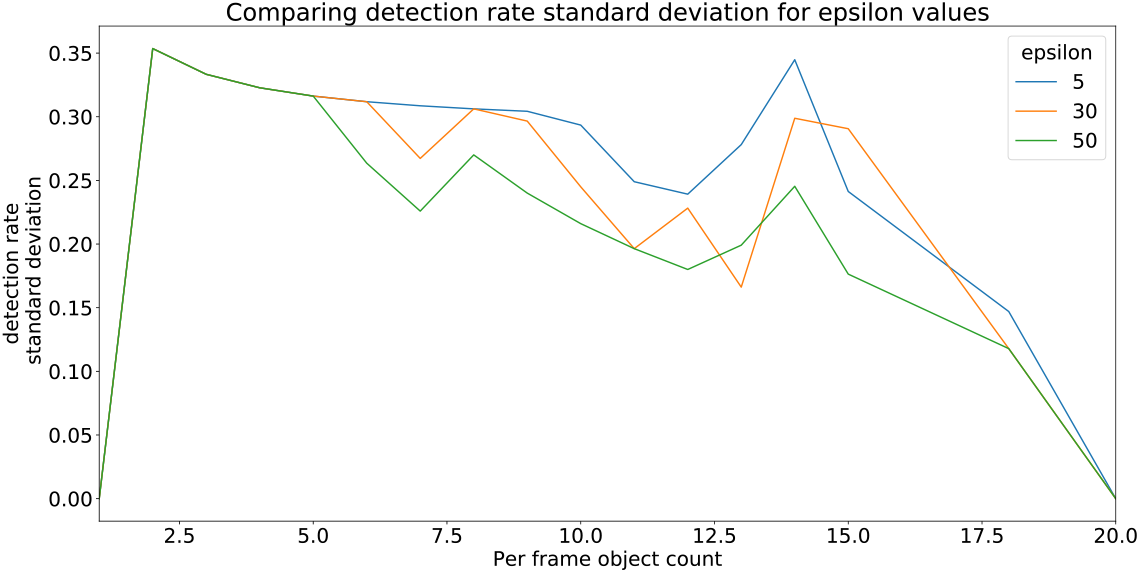
Standard deviation of detection rate for 200 frames in each of the three sample roots.

## 3 Methods

### 3.1 Data acquisition

To visualize mitotic events, we used the existing *Arabidopsis* transgenic line expressing a fusion between the cyclin protein CYCB1;1 and the fluorescent protein GFP, driven by the endogenous CYCB1;1 promoter [31]. The resulting fluorescent reporter CYCB1::GFP accumulates in cells transitioning between G2 and M phases of the cell cycle, and is quickly degraded after entering mitosis [31] and for this reasons it is widely adopted as a reliable live marker for mitotic events. *Arabidopsis* seeds were sterilised and stratified before sowing and germination on solid MS plates according to standard protocols [32]. Plants were imaged 5 days post-germination. For this study we imaged and analyzed *N* = 3 roots. A single primary root was grown and imaged on a custom-made light-sheet microscope setup, as previously described [16]. In essence, the root was hydroponically grown in a perfusion 5ml cuvette maintained under constant light and temperature, with its liquid medium fully exchanged every 2 minutes. A full 3D scan in fluorescence of the root tip generates a stack of 60 optical sections 4*μ*m apart, captured every 15 minutes for up to 7 days.

### 3.2 Pre-processing

#### 3.2.1 Noise analysis

Noise and histogram levels were used on normalised grey-scale images taking pixel values in [0,1]. The noise shown in the top panel of Fig. 8b was estimated using a wavelet-based estimator of the Gaussian noise standard deviation[33, 34]. The middle panel in Fig. 8b describes the signal. This is a simple quasi-signal metric defined as a ratio between the 99^*th*^ and 95^*th*^ histogram percentile boundary values, which are determined empirically for our data to separate background from signal. When this value approaches 0, the corresponding signal appears very low. For values above a calibrated threshold, the corresponding signal lies outside the 99^*th*^ percentile of the data, which typically happens for isolated bright points in the image. A noise range was used to guide downstream strategies in the image processing pipeline. An empirically determined “normal” noise band [0.01, 0.045] was used in the sample parameters discussed here. When the noise was below 0.01 no denoising was necessary, otherwise wavelet denoising was applied. Noise exceeding 0.045, typically corresponded to light saturation and/or low signal-to-noise ratio, and was due to a low number of mitotic events. In such instances additional thresholding was applied to the data before wavelet denoising. When the noise exceeded 0.1, the signal-to-noise ratio was so low that the frame was marked as degenerate. Fig. 8a contrasts different noise levels in sample images and Fig. 8b plots image properties over time for a given experiment.

#### 3.2.2 Region of interest

A 2D projection of the data was obtained by summing the 3D tensor along the z-axis. An adaptive threshold based on image histogram percentile ranges was used to construct a narrow-band filter for the 2D data. This range can vary erratically between frames as shown in the bottom time series in Fig. 8b. A thin slice of the image data range (shaded band) was selected and an aggressive “Gaussian smoothing” (averaging neighbouring pixel values using a Gaussian kernel) with an empirically-determined sigma value (*σ* = 8) was used to find a mesh-like connected component corresponding to the region of activity in the root tip. The 2D mask of this largest component was extended to a 3D mask by projecting the 2D region back into the z-plane. The largest connected component in the thresholded image was identified as the root tip. These stages are illustrated in Fig. 12. The image processing pipeline continues to process data only within the ROI.

**Fig 12:**
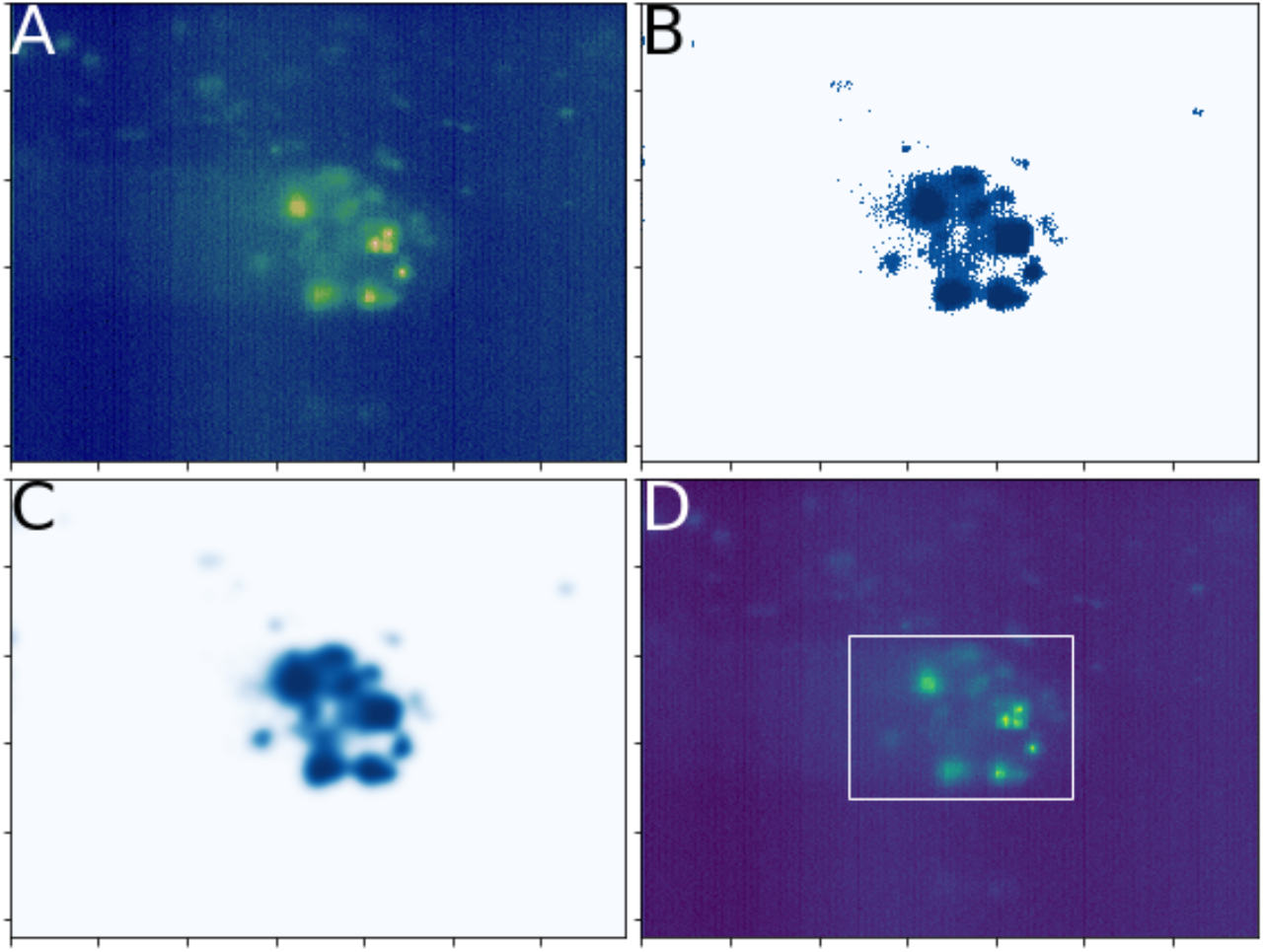
Isolating the Region of Interest (ROI): a narrow-band filter (B) and aggressive Gaussian smoothing (C) is used to find the largest connected component in the narrow band of the data corresponding to the region of activity in the root tip (D).

For cases where the region of activity is well-isolated (smaller region of interest), the processing is more efficient as the amount of volumetric data is reduced and debris beyond the root are filtered from the downstream pipeline. For our sample data, an ROI with area ≈ 10^5^ pixels corresponded to a well-isolated root tip. Noise and light saturation can affect this part of the process making the ROI area a proxy for image quality. While the actual area will depend on the data, it should vary smoothly and be relatively small unless root activity genuinely extends to the entire field of view.

#### 3.2.3 Object detection

We performed a “pre-segmentation” procedure to detect blob centroids. Segmentation plays a central role in many image processing pipelines and typically involves (i) thresholding and identifying background, (ii) using distance/gradient transformations with peak detection to identify markers and (iii) routines such as watershed [35] to segment blob labels. For our data, we have found it appropriate not to carry out the final segmentation. Instead, we carried out the pre-segmentation steps from thresholding to peak detection in detecting object centroids. Given the variability in the data over a large frame sequence, we have found a simple “annealing thresholding” to be effective. This simply increases a threshold iteratively so as to remove large connected components beyond a maximum perimeter length (empirically determined from data to be in the range 1000-2000 pixels) thus removing the background from the image. Being iterative, this is not as efficient as simpler linear filters but it is simple and robust to noise and variability. In extreme cases where it was not possible to find a threshold level in this manner, we treated the 99^*th*^ percentile as image background and removed it.

Having applied the threshold filter, we identified centroids by (i) performing a difference of Gaussians to emphasise blob-like objects, (ii) applying a maximum filter and (iii) returning the coordinates of the local maxima (peaks) in the image. To allow for a fully automated routine that can cope with arbitrary datasets (in the scope of our light-sheet microscopy datasets) the emphasis in the centroid detection stage has been to avoid spurious centroid detections at the risk of under-sampling, while optimising for objects to be identified for at least two frames somewhere during the peak of their light intensity arc.

### 3.3 Notes on algorithm assumptions

We consider the role of certain model assumptions in coping with variable and noisy data.

#### 3.3.1 Catchment region

Objects in the point cloud were expected to be separated by a minimum distance *ε* such that under global transformation single objects are matched.The parameter *ε* is of central importance at all stages of the image processing and tracking pipeline and can be chosen based on the data. For sparse point-cloud data there will typically be only one match within the ball of radius *ε*. An *ε* value can be chosen such that it is possible to find multiple objects matched to one object. The interpretation in this case is that all matched points in the catchment region correspond to the same object and that multiple points are the result of *e.g.* image processing anomalies. Merging lineages can be useful to avoid generating superfluous new identifiers. This adds robustness in the event of noisy data or image processing anomalies. Alternatives to using a ball of radius ε might consider anisotropic catchment regions that factor in the direction of the global transformation. The parameter ε value may result in the tracker merging objects that the object detection stage discriminates between if those objects are not sufficiently separated. In the case of accurately counting mitotic events, this reduces the introduction of false-positives *i.e.* the erroneous generation of new identifiers

#### 3.3.2 Consensus translation

Object position updates were treated as small Gaussian perturbations of a global transformation. If the position of an object is given as

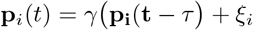

where 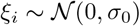 is the random fluctuation for one object, taking the “consensus” effectively filters out the noise to reveal the global movement. For example, the top *m* translations could be averaged to produce a new translation.

#### 3.3.3 Minimum allowed frame detections

Transient events were expected to persist for a minimum number of frames greater than or equal to 1. For our data the minimum number of frames is 2. Objects that have been identified for less than 2 frames, termed *single-frame detections*, were excluded from the tracking result.

#### 3.3.4 Gaussian distribution of ROI likelihood

In each frame, the objects of interest were expected to be found in the center of the field of view. Objects may be detected near the original frame boundaries depending on the size of the ROI. Objects can be treated as inliers/outliers based on standard outlier detection methods under the assumption of normality (for example within 1 or 2 standard deviations from the mean coordinate).

#### 3.3.5 Further filtering of outliers

Object location and angle differences between object displacements and the global transformation may be used in a post-processing stage to remove misclassified outliers. In the PRIORS routine in Algorithm listing 1 additional transformations beyond the transformations generated from the data can be chosen based on modelling assumptions. Additionally, the filter routines on lines 16-18 can filter data from the final result rather than during the tracking process based on modelling assumptions.

### 3.4 Software implementation

The fully automated Python software has been made available on GitHub at the link [36] and can be evaluated against test data available at the link [37]. The code is easily installed over a Python scientific library distribution such as *Anaconda* [38]. It includes interactive notebooks and can be run from a terminal window to process a folder containing 3D image sequence data in fully automated fashion.

CPU usage was dominated by the image processing stage. For the light-sheet microscopy experiments used in our evaluation, 3D images were approximately 90, 000KB on disk with dimensions (1392,1040, 60), in (*x, y, z*) order. The software was developed and tested on a personal laptop computer with 16GB RAM and a 2.9-GHz Intel Core i7-3520M CPU. While times vary between different pipeline modes, on average individual frame processing took around 40 seconds. More than half of the time spent on image processing is spent on de-noising and smoothing images. A negligible fraction of CPU time corresponds to the tracking stage of the pipeline. The tracking stage alone processed 450 data frames in about 1.5 minutes. The current implementation contains significant “meta analysis” overhead.

## 4 Discussion

In this work we address the problem of tracking transient events in structured 3D point clouds and in absence of background keypoints, a typical scenario experienced in extended *in vivo* time-lapse imaging of short-lived events in whole plant tissues. We present a new tracking method and validate it by tracking cell divisions in *Arabidopsis* three independent root meristems, over the course of several days. The novelty of the method proposed is in the way it deals at the same time with the transient nature of the events to be tracked, and the lack of background reference points, or keypoints. The validation presented against manual tracking of experimental data shows the improved performance of this method compared to previously published approaches on similar datasets.

Due to the transient nature of mitotic events and to the variation in image quality in experiments lasting up to one week, we found a lack of guaranteed, permanent features in image sequences. To circumvent the lack of fiducial markers, we introduced the concept of fuzzy markers defined through sub-sampling the set of detected events. In essence, we exchanged a morphological analysis problem for a fuzzy registration-tracking problem. The algorithm then generates and evaluates candidate affine transformations between these fuzzy markers. This approach is distinct from existing single-particle 3D tracking methods, which typically put greater emphasis on the existence of inlier objects. Furthermore, our method is characterised by a rigid-structure prior, since the algorithm models frame-frame object movement as a superposition of (a) long-range affine transformations due to global movement of the tissue and (b) short-range transformations due to local fluctuations of the nuclei within the tissue. The validation presented against both manual tracking and surrogate data indicates the validity of the method in tracking transient events in a 3D space.

It should be noted that the described algorithm has not been optimised for large point clouds. The point clouds we worked with contain less than 100 objects in each frame and often less than 20. As the frames are large 3D volumetric images, image processing is the performance bottleneck. Consequently, we have focused on the analysis and flexibility of the tracking algorithm instead of its computational optimisation. The transformation sampling and evaluation stages have been implemented efficiently within a vector-programming paradigm. Focus on performance improvements should emphasise efficiency of the *transformation proposal sampling*. This should, in general, be considered *NP-hard* as proposals are generated by finding congruences in large point cloud data. Also, for point clouds with high objects counts, proposal sampling could be applied on a suitable subregion of the data to avoid excessive evaluations.

## 5 Conclusion

In extended time-lapse microscopy, image processing and object detection relies heavily on checking and responding to variability in image properties. This goes beyond adaptive thresholding and may result in taking different routes through an image processing pipeline. Isolation of regions of interest within noisy point cloud data is also important.

Our approach was motivated by the case of transient florescent markers used in biology, where sub-cellular fluctuations are superposed with large-scale movements due to tissue growth. We present a clear case-study based on data on mitotic divisions in growing *Arabidopsis* roots, which could be immediately applied to similar datasets from other tissues. Given the vast variability in image quality and morphological characteristics (*e.g.* tissue types, density of objects to be tracked, landmark reference points, etc) in time-lapse microscopy datasets in life sciences and plant biology in particular, we believe that the novel approach proposed here will be a useful addition to the existing toolbox for single-particle tracking in 3D. Moreover, we expect that our approach will be applicable to any other situations where tracking transient events embedded in rigid structures is required.

## Declarations

### Author’s contributions

SA developed the code, carried out most of the data analysis and wrote most of the manuscript; TF carried out the microscopy experiments and contributed to the data analysis; GP contributed to the code development; GS conceived and designed the project and contributed to the writing of the manuscript; all authors reviewed the manuscript.

### Availability of data and materials

The Python code [36] can be run against sample data hosted in a publicly accessible data repository [37]. The free-to-use code, which has an MIT license, can be run on any operating system running Python. The README at the project homepage provides instructions to run the code with the sample data.

### Ethics approval and consent to participate

Not applicable

### Consent for publication

Not applicable

## Acknowledgements

Not applicable

## Supplementary Figures

